# miRNAs play important roles in aroma weakening during the shelf life of ‘Nanguo’ pear after cold storage

**DOI:** 10.1101/247932

**Authors:** Fei Shi, Xin Zhou, Miao-miao Yao, Zhuo Tan, Qian Zhou, Lei Zhang, Shu-juan Ji

**Author notes:** These authors contributed equality to this study (Fei Shi, Xin Zhou). Corresponding author: Dr Shujuan Ji, Department of Food Science, Shenyang Agricultural University, No. 120 Dongling Road, Shenyang 110866, P.R. China. *E-mail. **Abbreviations:** RT: room temperature LT: low temperature OTP: optimum tasting period SSC: soluble solid content miRNAs: microRNAs LOX: lipoxygenase HPL: hydroperoxide lyase ADH: alcohol dehydrogenase AAT: alcohol acyltransferase FC: fold change RT-qPCR: reverse transcription quantitative PCR.

## Abstract

Cold storage is commonly employed to delay senescence in ‘Nanguo’ pears after harvest. However, this technique also causes fruit aroma weakening. MicroRNAs play important roles in plant development and in eliciting responses to abiotic environmental stressors. In this study, the miRNA transcript profile of the fruit at the first day (C0, LT0) move in and out of cold storage and the optimum tasting period (COTP, LTOTP) during shelf life at room temperature were analyzed, respectively. More than 300 known miRNAs were identified in ‘Nanguo’ pears; 176 and 135 miRNAs were significantly differentially expressed on the C0 *vs*. LT0 and on the COTP vs. LTOTP, respectively. After prediction the target genes of these miRNAs, *LOX2S*, *LOX1_5*, *HPL*, and *ADH1* were found differentially expressed, which were the key genes during aroma formation. The expression pattern of these target genes and the related miRNAs were identified by RT-PCR. Mdm-miR172a-h, mdm-miR159a/b/c, mdm-miR160a-e, mdm-miR395a-i, mdm/ppe-miR399a, mdm/ppe-miR535a/b, and mdm-miR7120a/b negatively regulated target gene expression. These results indicate that miRNAs play key roles in aroma weakening in refrigerated ‘Nanguo’ pear and provide valuable information for studying the molecular mechanisms of miRNAs in the aroma weakening of fruits due to cold storage.

## Introduction

‘Nanguo’ pear (*Pyrus ussuriensis* Maxim.), which is mainly produced in Liaoning Province, China, has soft flesh, a rich aroma, dietary fiber, sugar, protein, and various other nutrients when ripe. Its fruits are usually consumed fresh and are believed to provide health benefits. During the optimal taste period (OTP) of the fruit at room temperature (RT, 20°C ± 1°C), the pear has soft and smooth flesh, a sour-sweet taste, and an attractive aroma. Together with fruit color, shape, total soluble solid content, and titratable acidity, aroma is an important factor affecting fruit quality and final consumer acceptance (Infante et al., 2008). The aroma of ‘Nanguo’ pear is mainly due to aroma esters (Ji et al., 2012). The fruit ripens and rapidly deteriorates at RT after harvest and thus has a short shelf life. Low temperature (LT, 0°C ± 0.5°C) is an effective method for prolonging storage life by slowing fruit ripening and reducing decay. However, fruit aroma also weakens with cold storage, which in turn affects its quality and commodity value, especially following long-term LT storage (Zhou et al., 2014; Shi et al., 2017).

MicroRNAs (miRNAs) are a class of endogenous non-coding RNAs (18–24 nt in length) that mainly negatively regulate gene expression at the post-transcriptional level by binding to complementary sequences within target mRNAs (Jones-Rhoades et al., 2006; Mallory and Vaucheret, 2006; Meyers et al., 2008). MiRNA binding leads to either cleavage-induced degradation of the mRNA or suppression of its translation. In plants, the complementarity between miRNAs and their targets is very high, and miRNAs play critical roles in the regulation of plant growth (Mallory et al., 2004), development (Chen, 2004), and responses to abiotic stresses such as high salinity (Liu et al., 2008), high or low temperature (Zhang et al., 2009; Lv et al., 2010). LT storage is known as a major abiotic stress that affects plant development and growth and impacts aroma production (Josine et al., 2011; Zhou et al., 2014).

High-throughput degradome library sequencing has provided a new strategy for validating splicing targets at the whole-genome level and has been successfully employed in identifying various miRNAs and their target genes in plants (Addo-Quaye et al., 2008; Bartel, 2004a; Song et al., 2016; Xu et al., 2012, 2013). High-throughput sequencing has identified numerous critical miRNAs, such as miR156, miR159, miR160, miR164, miR172, miR319, miR393, miR394, miR395, miR399, and miR408, that are expressed in response to abiotic stresses in plants (Jones-Rhoades et al., 2006; Schommer et al., 2008; Stief et al., 2014; Sun, et al., 2015; Zhang et al., 2016). MiR156/157 negatively regulates the MADS-RIN to induce the transcription of ripening-related genes in tomato (Dalmay 2010), and the known ripening regulators, the CNR and AP2 TFs, were also identified as targets of miR156 and miR172, respectively (Chung et al., 2008; Karimi et al., 2016). In addition, miRNAs negatively regulate their targets in *Arabidopsis* roots after infection with cyst nematodes, leading to the downregulation of miRNAs (Xing et al., 2010). In *Arabidopsis thaliana*, *Populus trichocarpa*, and other plant species, miR156, miR159, miR172, and miR394 exhibit altered expression under cold stress (Liu et al., 2008; Lu et al., 2008; Zhou et al., 2008). To date, more than 30,000 miRNAs have been identified, but the function of most miRNAs remain unclear; miRNAs have not been reported in ‘Nanguo’ pear and their functions in fruit aroma weakening after cold storage remain unknown.

In our previous study, to understand the biological functions of miRNAs and to investigate the mechanism of fruit aroma weakening after LT storage, 12 mRNA libraries of ‘Nanguo’ pear at RT and after storage were constructed and sequenced on an Illumina HiSeq 4000 platform, and numerous mRNAs and TFs involved in aroma formation were identified. In the present study, to understand the role of miRNAs on fruit aroma weakening in response to cold storage, 12 miRNA libraries (at 0 d and OTP at RT and after LT storage, respectively, with three biological replicates each) from ‘Nanguo’ pears were constructed and sequenced on an Illumina Hiseq 4000 platform. Compared to the RT fruit, 176 miRNAs and 135 miRNAs were differentially expressed in refrigerated fruit at 0 d and at OTP, respectively. Prediction of the target genes of the differentially expressed miRNAs showed that these are related to aroma weakening after LT storage, and their possible roles were analyzed.

## Materials and Methods

### Plant materials and treatment

‘Nanguo’ pears were harvested at a commercial orchard in Anshan, Liaoning Province, China and transported to the laboratory on the day of harvest. Fruits without visible signs of defects or decay were selected and randomly divided into two groups (N = 400 fruits in each group), and stored at RT for 5 d premature. The fruits were subpackaged in 0.04-mm thick polyethylene (PE) bags to maintain a relative humidity of about 80%–85%, and then one group was stored at 20°C and monitored at time points 0 d, 3 d, 6 d, 9 d, and 12 d. The other group was stored at 0°C; the bag was opened for precooling 24 h, then the bag was and stored for 150 d, and subsequently transferred to RT and monitored at time points 0 d, 3 d, 6 d, 9 d, and 12 d. The fruit samples on the first day (C0, LT0) move in and out cold storage, and on the OTP (COTP, LTOTP) during shelf life were as the sequencing samples. The test comprised three replicates of 30 fruits for each treatment. The fruit peels of each sample at various time points were immediately frozen in liquid nitrogen and stored at −80°C.

### Evaluation of the fruit ripening

Fruit firmness was measured with a texture analyzer using a 2-mm plunger tip (TA-XT2i Plus, Stable Micro System, UK). The test rate was 3 mm·s^−1^ to a 5-mm depth on opposite sides of the equator of 18 fruits after removal of a slice of skin. Ethylene was collected from the box headspace and analyzed using a gas chromatograph (CP-3800, Varian, USA) equipped with a flame ionization detector (the injector, detector, and oven temperatures were 110°C, 140°C, and 90°C, respectively). Four fruits from each treatment were weighed and placed in the 1.2-L fresh-keeping box for 5 h at RT. Soluble solid content (SSC) were measured using a digital hand-held refractometer (PAL-13810, Atago, Tokyo, Japan) and expressed as percentage (%). The test was performed using the flesh of 18 fruits from the two opposing sides.

### miRNA Illumina sequencing and library construction

MiRNAs was extracted from fruit samples on the day move in and out of the cold storage (C01-03, LT01-03) and at OTP (COTP1-3, LTOTP1-3), with three biological replicates. Extraction was performed using an miRNA isolation kit (CWBIO, Beijing, China) following the instructions provided by the manufacturer. RNA quality was monitored by gel electrophoresis and an OD_260_/OD_280_ ratio. High-quality RNA samples were used for miRNA library preparation and massive sequenced by an HiSeq 4000 SBS kit (300 cycles) (Illumina, San Diego, CA, USA). Multiplexing was applied during sequencing. Libraries were constructed according to TruSeq^TM^ Small RNA sample prep kit (Illumina, USA). Briefly, total RNA was purified by electrophoretic separation on a 15% TBE-urea denaturing polyacrylamide gel electrophoresis (PAGE) gel, and miRNA regions corresponding to the 18–32 nt bands in the marker lane were excised and recovered. Then, the18–32 nt miRNAs were ligated to a 5’-adaptor and a 3’-adaptor sequentially (TruSeq^TM^ Small RNA sample prep kit). The adapter-ligated miRNAs were subsequently transcribed into cDNA and then PCR amplified for 12 cycles using the adaptor primers. The PCR products were purified for high-throughput sequencing.

### Analysis of miRNA sequencing data

The raw data were assessed in terms of quality and base count. Then, the raw data were extracted from unknown and low-quality base sequences and then screened in terms of length. The remaining data were then designated as clean, small RNA sequences, and their length distribution and base preference were analyzed using *Fastx-Toolkit* (http://hannonlab.cshl.edu/fastx_toolkit). To analyze the conserved miRNAs, unique small RNAs were aligned with mature plant miRNAs from miRBase. Using the Rfam (12.1, http://Rfam.sanger.ac.uk/) database to annotate the measured small RNAs,the non-miRNA sequences were identified using BLASTn (http://blast.ncbi.nlm.nih.gov/). The miRNA sequences were compared to reference genomes using Bowtie (http://bowtie-bio.sourceforge.net/index.shtml). Small maize RNA sequences with one mismatched bases were identified as known plant miRNAs.

### Identification of differently expressed miRNAs and prediction of their target genes

Differentially expressed miRNAs were analyzed using the software DESeq2. Transcripts per million (TPM) reads were used to evaluate the relative expression levels of each miRNA in four small RNA libraries. If the read count of a given miRNA was zero, then the TPM was modified to 0.01 and excluded from analysis of differential expression. Differences in fold-change among the four libraries were calculated using the following equation: Fold-change = log_2_ (FC). The miRNAs with FCs > 1 or < −1 and *p*-values ≤ 0.05 were respectively considered upregulated or downregulated in response to low temperature.

Target gene predictions were performed based on the criteria described by Enright et al. (2003). Previous reports have shown that miRNAs are capable of binding to both coding sequences (CDs) and the 3’ untranslated regions (3’UTRs) of a transcript, regulating the expression of the target gene (Bartel, 2004b; Ren and Yu, 2012).

### RT-PCR analysis of differentially expressed miRNAs and their target genes

‘Nanguo’ pears were sampled on 0 d, 3 d, 6 d, 9 d, and 12 d of shelving at room temperature and after cold storage for RT-PCR of miRNAs and their target genes. MiRNA and RNA were extracted using an miRNA isolation and OminiPlant RNA kit according to the manufacturer’s instructions. The miRNAs and RNA were subsequently converted into cDNA using an miRNA and RNA cDNA synthesis kit (CWBIO, Beijing, China), which was then used as template for RT-PCR. RT-PCR amplification and analysis were performed using QuantStudio 5 Flex (Life Technologies, USA) with a miRNA and RNA RT-PCR assay kit. The RT-PCR conditions were based on the instructions of the RealMasterMix kit (SYBRGreen, CWBIO, Beijing, China). The levels of 166miRNA and target genes were calculated using the 2^ΔΔ^Ct method, with the actin gene (PuActin, JN684184) used as internal control (Zhou et al., 2014). Three biological repeats were used in the RT-PCR assay. The primer sequences used in this study are shown in Table 4.

### Statistical analysis

All data were analyzed by one-way ANOVA. Mean separations were performed by Duncan’s multiple range test at the 5% level, standard errors (±SE) were calculated using Microsoft Excel (3 replicates; Redmond, WA, USA).

## Results

### Maturity of ‘Nanguo’ pears during postharvest storage

Fruit maturity was evaluated by measuring the firmness, ethylene production, and SSC during shelf life at RT and after cold storage, as well as changes in fruit aroma relative to stages of maturation (Ji et al., 2012). Table 1 shows that the firmness values of ‘Nanguo’ pears after release from LT storage for 150 d were significantly lower than those recorded on the day before cold storage was initiated, and firmness rapidly decrease to below that at RT, which both decreased during shelving. A 6.5-7.5 N firmness value was considered indicative of moderate softness and the optimal time for tasting ‘Nanguo’ pears (Zhou et al., 2014; Shi e al., 2017). Ethylene production was significantly higher after cold storage for 150 d (*P* < 0.05); it peaked on the sixth day, which was three days earlier than that observed in fruits stored at RT. No changes in TSS during shelving were observed in the two fruit groups, which initially increased, peaking on at 9 d and 6 d at RT and LT, respectively, and then subsequently decreased. The results indicated that rate of fruit ripening increased after cold storage, and the OTP was on the 9 d and 6 d in fruits stored at RT and LT, respectively.

**Table 1.**
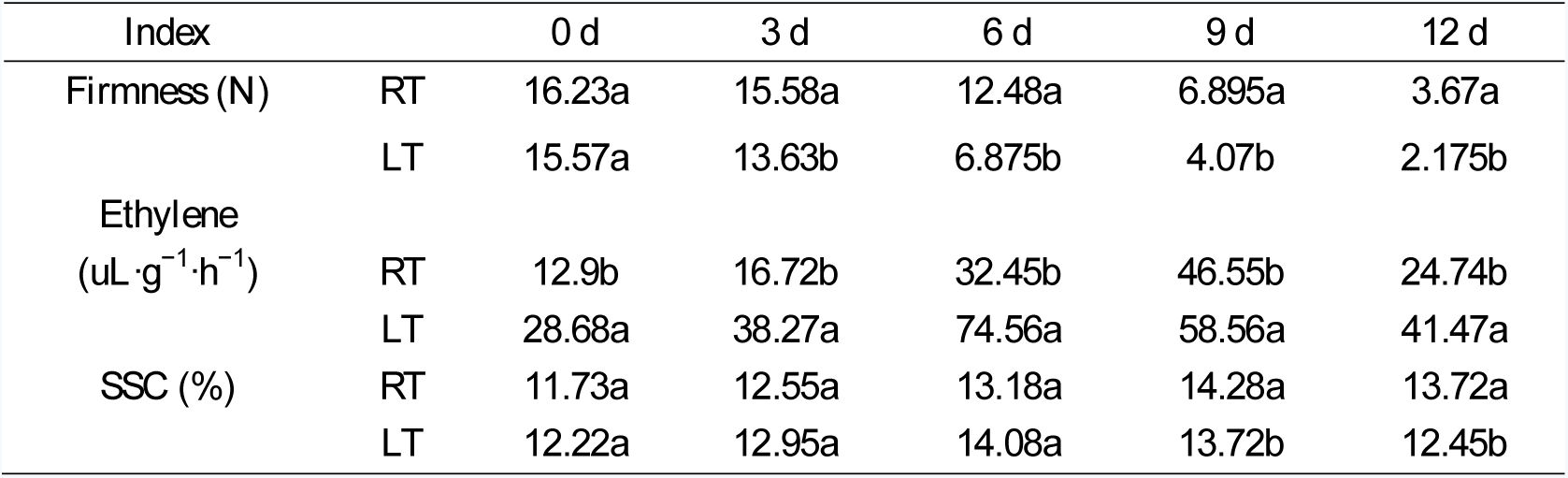
Changes of firmness, ethylene and soluble solid content (SSC) during shelf life of ‘Nanguo’ pears at room temperature (RT) and after 150 d low temperature storage (LT).

### Sample collection and overview of the miRNA sequencing of ‘Nanguo’ pears

To investigate the function of miRNAs in aroma weakening in ‘Nanguo’ pears during cold storage, fruits stored at RT and LT were sampled on the first day and at OTP, respectively. Twelve libraries (C01-03, COTP1-3, LT01-03, and LTOTP1-3) of small RNAs were constructed for the samples by Illumina sequencing. Table 2 shows that an average of 16,490,952 raw reads were obtained from the 12 libraries of the small RNA sequence of ‘Nanguo’ pears. After quality control of the raw reads, 7.95-19.4 million clean-read distinct sequences were obtained, with sizes ranging from 18 nt to 32 nt; with two major peaks at 21 nt and 24 nt (Figure 1A). The number of reads for the 21-nt sequences was higher in the RT samples; however, the reads of the 24-nt was higher in the LT samples. As shown in other plants, the 21-nt and 24-nt miRNAs may play a role in maintaining genome integrity and stabilization via heterochromatin formation.

**Table 2.** Summary of the raw date of all microRNAs in the room temperature fruit (C0 and COTP) and the cold storage fruit (LT0 and LTOTP).

| Sample-ID | Total Reads | Total Bases | Error% | Q20% | Q30% | GC% | Clean reads |
| --- | --- | --- | --- | --- | --- | --- | --- |
| C01 | 13303737 | 678490587 | 0.0108 | 99.3 | 97.89 | 51.74 | 15886825 |
| C02 | 13070747 | 666608097 | 0.0109 | 99.3 | 97.87 | 51.83 | 13962727 |
| C03 | 11614705 | 592349955 | 0.0103 | 99.56 | 98.46 | 52.34 | 7953644 |
| COTP1 | 15036061 | 766839111 | 0.0107 | 99.38 | 98.08 | 52.23 | 14197822 |
| COTP2 | 12337321 | 629203371 | 0.0103 | 99.57 | 98.47 | 52.86 | 19403640 |
| COTP3 | 12040228 | 614051628 | 0.0103 | 99.57 | 98.46 | 53.43 | 17574539 |
| LT01 | 18091065 | 976917510 | 0.0099 | 99.37 | 97.77 | 50.29 | 12454217 |
| LT02 | 25532143 | 1378735722 | 0.0099 | 99.36 | 97.73 | 49.75 | 9974659 |
| LT03 | 23318692 | 1259209368 | 0.01 | 99.3 | 97.61 | 49.87 | 9597669 |
| LTOTP1 | 22473351 | 1213560954 | 0.01 | 99.33 | 97.65 | 50.12 | 11915780 |
| LTOTP2 | 20576979 | 1111156866 | 0.0101 | 99.25 | 97.45 | 49.87 | 11503596 |
| LTOTP3 | 10496389 | 566805006 | 0.01 | 99.31 | 97.61 | 50.02 | 9216281 |

**Fig. 1.**
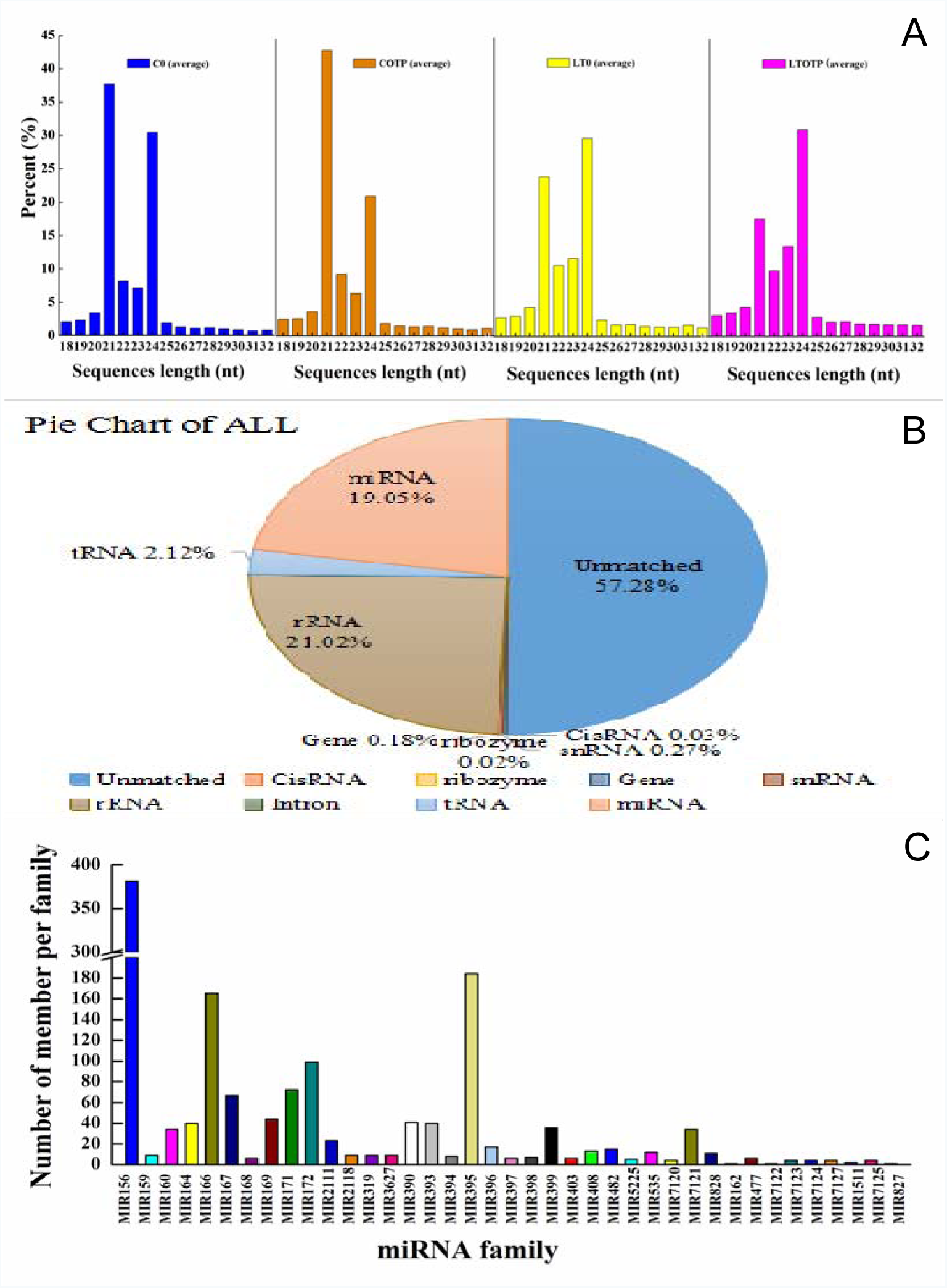
A: Length distribution of sRNA reads and unique sequences in the room temperature fruit (RT) and low temperature storage fruit (LT) twelve libraries constructed from ‘Nanguo’ pears. B: Comparison the unique sequence of all samples to the Rfam database. C: Distribution of known miRNA family size in ‘Nanguo’ pears.

### Identification of known miRNAs

After removing the rRNAs, snRNAs, tRNAs, and non-miRNA sequences, approximately 656,407,32 reads were matched in the all 153,641,399 reads, and 292,720,11 reads (19.05%) were matched to miRNAs (Figure 1B). To identify known miRNAs in the 12 libraries of ‘Nanguo’ pears, these sequences were mapped to the genome of *Prunus persica* and *Malus domestica* using bowtie, allowing one mismatch with less than two substitutions to known miRNAs from miRBase. By querying the remaining sequences against the currently known plant precursor or mature miRNA sequences in miRBase21 (http://www.mirbase.org/), more than 300 known miRNAs belonging to 46 miRNA families were identified in ‘Nanguo’ pear that mainly matched those of *M. domestica* and *P. persica*. Comparison of family sizes showed that the number of miRNA members widely varied among miRNA families. The number of members in each miRNA family in ‘Nanguo’ pear ranged from 1 to 380, with families miR156, miR166, MIR172, and miR395 showing the highest number of members (Figure 1C).

### Identification of the differentially expressed miRNAs in response to LT storage

To identify differentially expressed miRNAs related to aroma formation during cold storage, the miRNAs identified in the LT0 and LTOTP groups were compared to those of the C0 and COTP groups, whereas those of the C0 and COTP groups were used as control. The number of differentially expressed miRNAs of the CO *vs*. COTP and LT0 *vs.* LTOTP ‘Nanguo’ pears were analyzed, using those from the C0 and LT0 groups as control. Figure 2B shows that 12 miRNAs, including mdm-miR396g/f, mdm-miR397a/b, mdm-miR398b/c, ppe-miR398b/a-3p, and mdm-miR408b/c/d were downregulated, and 14 miRNAs consisting of ppe/mdm-miR858, mdm-miR168a/b,mdm-miR156t/u/v/w,ppe/mdm-miR394a/b,and mdm-miR827were upregulated between C0 and COTP. However, 36 miRNAs, including mdm-miR156a-o, ppe-miR156c/d/e, mdm-miR1511, mdm-miR172a-h, ppe-miR169f-j, and ppe/mdm-miR858 were upregulated, and only four miRNAs, including mdm-miR399a/b/c and ppe-miR399b, were downregulated in LT0 *vs*. LTOTP (Figure 2C). These results show that miRNAs are significantly differentially expressed during aroma formation before and after cold storage. Compared to the miRNA expression in fruits stored at RT,117 miRNAs, including mdm-miR 167b-g, mdm-miR162a/b, mdm-miR172a-o, ppe/mdm-miR394a/b, mdm-miR399a/b/c, mdm-miR164b-f, mdm-miR160a-e, mdm-miR7120a/b, mdm-miR7121a/b/c, mdm-miR159a/b, mdm-miR156t/u/v/w, 229mdm-miR396b,mdm/ppe-miR395a-f,mdm-iR171a/b/f/h,mdm-miR167i/j,and mdm/ppe-miR535a were upregulated, and 59 miRNAs, including ppe/mdm-miR166a-i, 231mdm-miR482a-5p,ppe-miR398b,mdm-miR171i,mdm-miR156a-o,mdm-miR398b/c, mdm-miR397a/b, mdm-miR408a/b/c/d, and mdm-miR159c were downregulated between C0 and LT0 (Figure 2D), and 108 miRNAs, including ppe-miR398b, mdm-miR172a-o, mdm-miR167b-g, 234mdm-miR156t/u/v/w,mdm/ppe-miR394a/,mdm-miR159a/b,mdm/ppe-miR395b-i, mdm-miR160a-e, mdm-miR7120a/b, mdm/ppe-miR535a, mdm-miR164b-f, mdm-miR2118a/b/c, and mdm-miR168a/b were upregulated, and 27 miRNAs consisting of ppe/mdm-miR166a-i, mdm-miR408a, mdm-miR393a/b/c, mdm-miR7121d-g, mdm-miR398b/c, and mdm-miR319a/b were downregulated between COTP and LTOTP (Figure 2E). These findings show that the miRNA expression is significantly altered during cold storage, particularly mdm-miR172a-o, mdm-miR167b-g, mdm/ppe-miR395a-i, mdm-miR160a-e, mdm-miR7120a/b, mdm-miR159, mdm-miR164b-f, and mdm-miR396b, which were only significantly upregulated after LT storage.

**Fig. 2.**
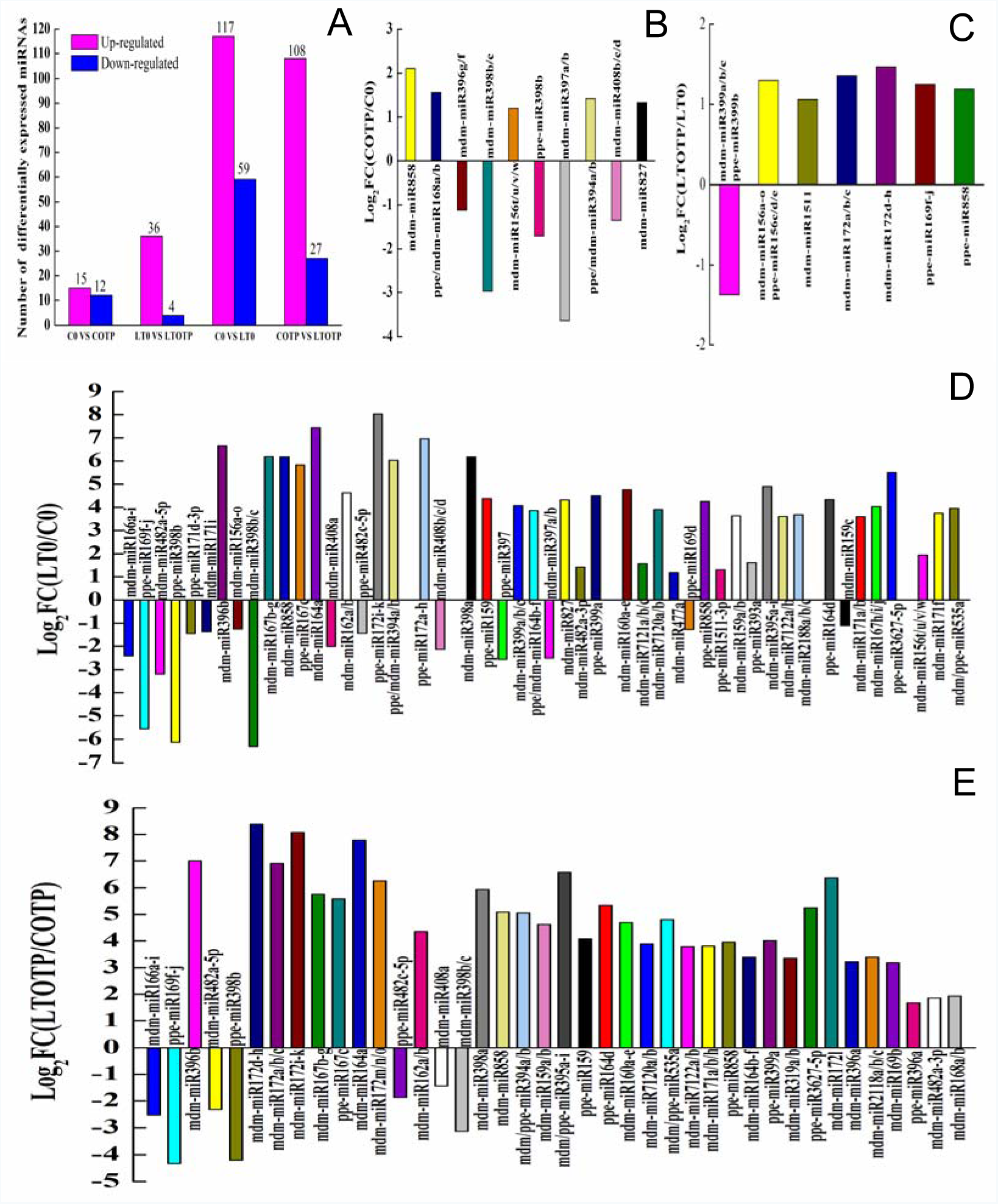
The number of up-regulated and down-regulated miRNAs between the four compared groups is summarized in A, and the expression pattern of the significantly differentially expressed miRNAs in different compared libraries are shown in B, C, D and E. B: C0 VS COTP, C: LT0 VS LTOTP, D: C0 VS LT0, E: COTP VS LTOTP.

### Identification of the target genes of the differentially expressed miRNAs

MiRNAs regulate their target genes, some miRNAs may have multiple target genes, and the same target gene may also be regulated by multiple miRNAs. Several target genes of the differentially expressed miRNAs were identified, and some of the target genes were key enzyme genes of aroma formation such as *LOX2S*, *LOX1_5*, *HPL*, and *ADH1*. As shown in Figure 3, *LOX2S* may be regulatedbymdm-miR535a/b,mdm-miR160a/b/c,mdm-miR156a-o/t-w,mdm-miR408a-d, mdm-miR159a/b/c,mdm-miR168a/b,mdm-miR172a-o,mdm-miR394a/b,mdm-miR395a-h, mdm-miR396a/c/e, and mdm-miR7120a/b. *LOX1_5* may be regulated by mdm-miR408b/c/d, mdm-miR159a/c,mdm-miR399a/b/d,mdm-miR7120a/b,mdm-miR169b,and mdm-miR172c/e/f/g/h. *HPL* is regulated by mdm-miR7120a/b. *ADH1* may be mainly regulated by mdm-miR394a/b, mdm-miR395a-h, mdm-miR827, mdm-miR156a-o/t-w, mdm-miR168a/b, and mdm-miR408b/c/d. The four target genes are involved in the fatty acid metabolism pathway, and are key genes in aroma formation. Other target genes included signal transduction genes in the plant hormone signal transduction pathway and transcription factors in response to chilling injury. Correlation analysis indicated that the *LOX2S* gene is mainly regulated by mdm-miR160a-e, mdm-miR172a-h, mdm-miR535a/b, mdm-miR394a/b, mdm-miR395a-i, and mdm-miR159a/b/c; the *LOX1_5* gene is mainly regulated by mdm/ppe-miR399a and mdm-miR159a/b/c; and the *ADH1* gene is mainly regulated by mdm-miR156a-o (Tables 3a and b).

**Table 3.** The correlation coefficients between the change of the significantly differentially expressed miRNAs and the change of their predicted target genes LOX2S, LOX1_5, HPL, ADH1 on the 0 d and the OTP in ‘Nanguo’ pears after low temperature storage.

| C0 VS LT0 | LOX2S | LOX1_5 | HPL | ADH1 |
| --- | --- | --- | --- | --- |
| mdm-miR160a-e | -0.852* | - | - | - |
| mdm-miR172a-h | -0.833* | -0.768 | - | - |
| mdm-miR535a/b | -0.812* | -0.761 | - | - |
| mdm-miR394a/b | -0.842* | - | - | 0.907** |
| mdm-miR159a/b/c | -0.883* | -0.834* | - | 0.988** |
| mdm-miR399a/b/c | - | -0.820* | - | - |
| mdm-miR7120a/b | -0.823* | -0.753 | -0.677 | - |
| mdm-miR395a-i | -0.673 | -0.623 | - | 0.817* |
| mdm-miR408b/c/d | 0.468 | 0.483 | - | -0.776 |
| mdm-miR156a-o | 0.711 | - | - | -0.916** |

b
| COTP VS LTOTP | LOX2S | LOX1_5 | HPL | ADH1 |
| --- | --- | --- | --- | --- |
| mdm-miR160a-e | -0.955** | - | - | - |
| mdm-miR172a-h | -0.881* | -0.674 | - | - |
| mdm-miR535a | -0.842* | -0.712 | - | - |
| mdm-miR394a/b | -0.979** | - | - | 0.632 |
| mdm-miR159a/b/c | -0.931** | -0.699 | - | 0.676 |
| mdm-miR399a/b/c | - | -0.705 | - | - |
| mdm-miR7120a/b | -0.912* | -0.677 | -0.862* | - |
| mdm-miR395a-i | -0.980** | -0.671 | - | 0.766 |
| mdm-miR408b/c/d | 0.573 | 0.348 | - | -0.694 |
| mdm-miR156a-o | 0.673 | - | - | -0.816* |

\*, Correlation is significant at the 0.05 level, \*\*, Correlation is significant at the 0.01 level.

**Table 4.**
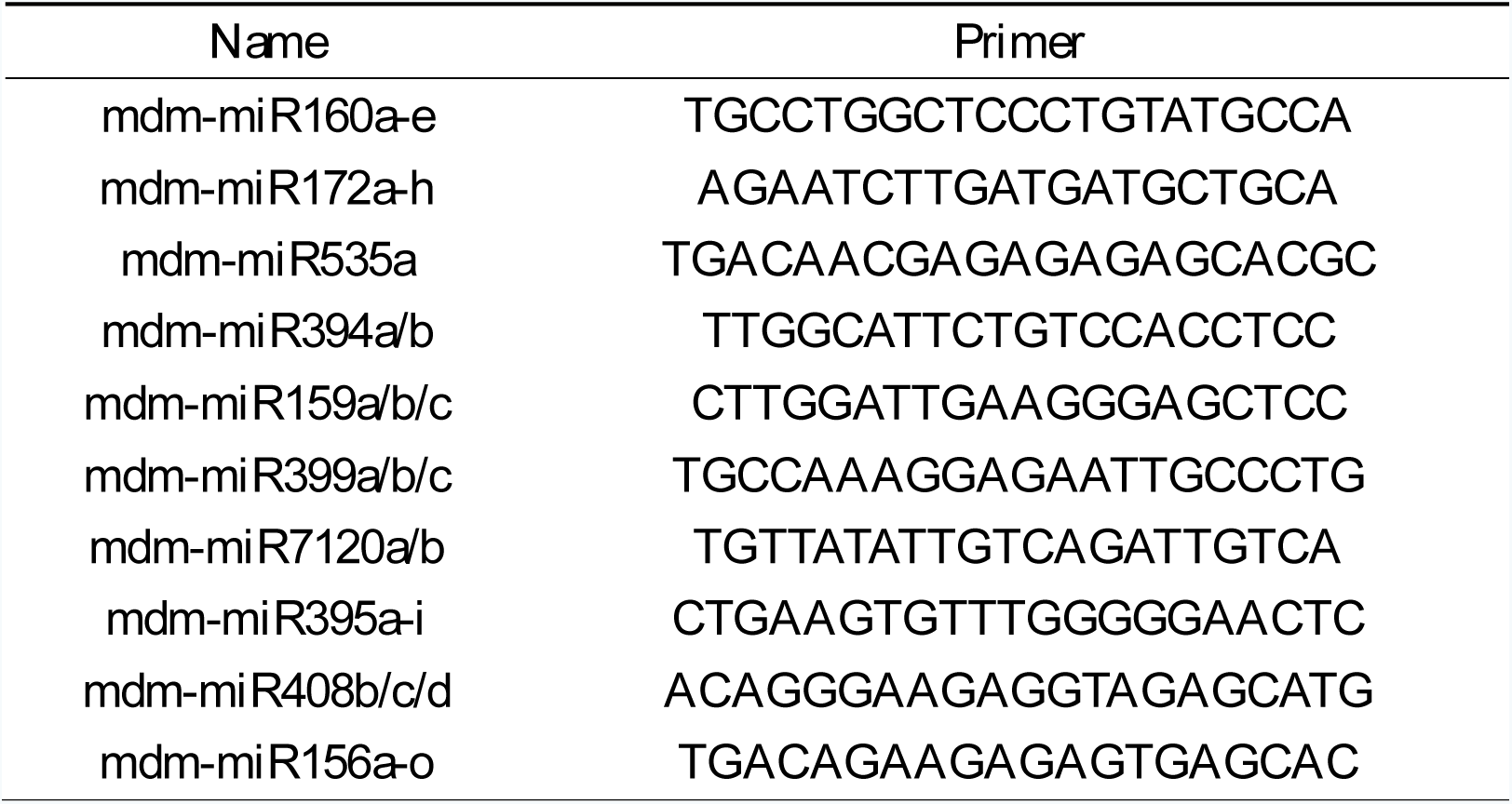
The primers of miRNAs for RT-PCR analysis.

**Fig. 3.**
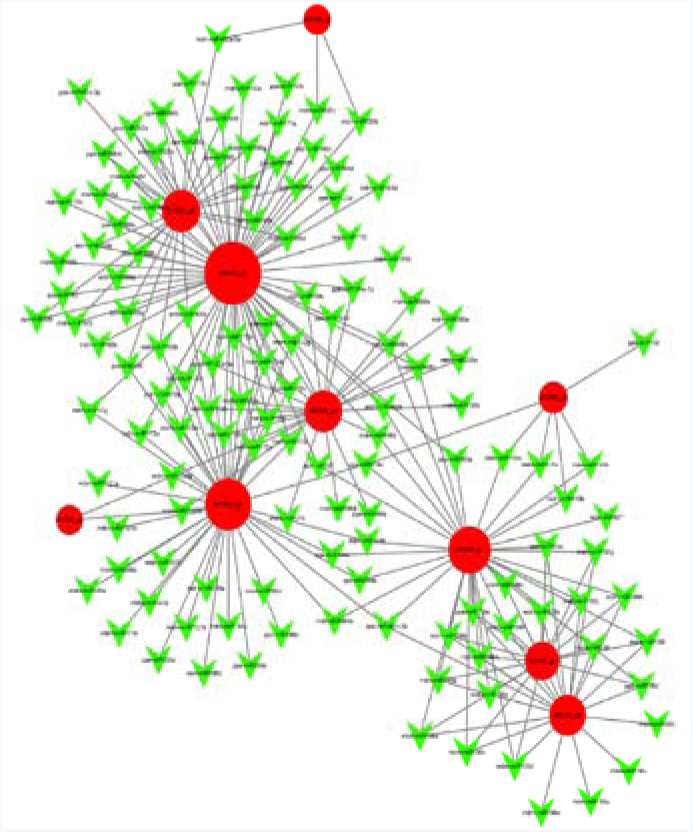
The network map of the significantly differentially expressed miRNAs and the target genes LOX2S, LOX1_5, HPL and ADH1. Red is the targer genes, green is the miRNAs. LOX2S:c62400_g2, c63403_g2, c65860_g1, c65968_g1, c66100_g1; LOX1_5: c54748_g2, c63599_g3; HPL: c64295_g2; ADH1: c27994_g3, c66564_g1.

### RT-PCR expression profiling of differentially expressed miRNAs and their target genes

To understand the role of various miRNAs in aroma weakening in ‘Nanguo’ pears during cold storage, the expression pattern of key genes related to aroma formation, namely, *LOX2S*, *LOX1_5*, *HPL*, and *ADH1*, were analyzed by RT-PCR (Figure 4A, 4B, 4C, 4D), simultaneous with those of miRNAsmdm-miR7120a/b,mdm-miR172a-h,mdm/ppe-miR394a/b,mdm-miR395a-h, mdm-miR535a, mdm-miR160a/b/c, mdm-miR156a-o, mdm-miR408a-d, mdm-miR159a-c, and mdm/ppe-miR399a. Figure 4 shows that compared to fruits stored at RT, the *LOX2S* and *HPL* genes were significantly downregulated on the day these were moved out of cold storage, and the *LOX2S* and *LOX1_5* genes were significantly downregulated at the OTP. And mdm-miR172a-o, mdm-miR395a-h,mdm/ppe-miR394a/b,mdm/ppe-miR399a,mdm-miR159a-c, mdm-miR160a/b/c, mdm-miR7120a/b and mdm-miR535a were significantly upregulated, and mdm-miR408a-d was significantly downregulated in the fruits compared to that stored at RT. MiRNAs negatively regulate the expression of target genes; therefore, mdm-miR172a-h, mdm-miR159a/b/c,mdm-miR160a-e,mdm-miR395a-h,mdm/ppe-miR399a, mdm/ppe-miR535a/b, and mdm-miR7120a/b may play important roles in aroma weakening in ‘Nanguo’ pears after cold storage by regulating these target genes.

**Fig. 4.**
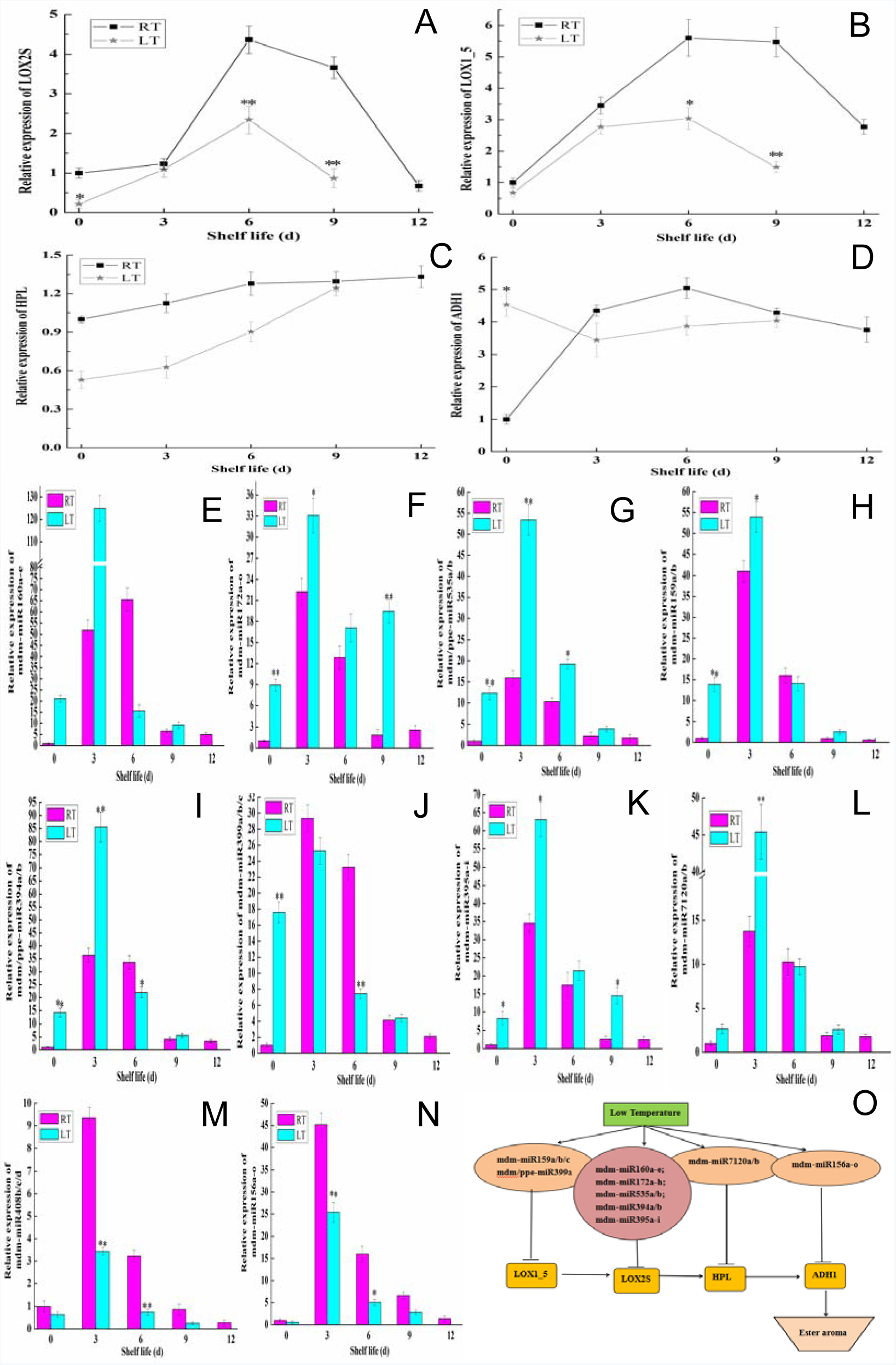
Expression analysis of significantly differentially expressed miRNAs and their predicted target genes in ‘Nanguo’ pear by RT-PCR. The symbols * and ** in the graph show the significant differences at *p* < 0.05 and *p* < 0.01, respectively.

### Discussion

‘Nanguo’ pears are characterized by aroma esters, which influence their commercial value. Studies have shown that maturity and senescence are critical factors affecting the abundance of aroma compounds in fruits (Lara, 2003), and that these processes induce the synthesis and degradation of aroma volatiles (Manriquez et al., 2007; Tietel et al., 2011). ‘Nanguo’ pears evolve through the stages of ripening, full ripening, and overripening during postharvest RT and LT storage. To ensure consistency in maturation stage at OTP before and after cold storage, the present study assessed fruit maturity under various storage conditions in relation to maturation. Here, firmness, ethylene production, and SSC were used as indices of fruit maturity during storage at room temperature and after cold storage. The results showed that fruit firmness was lower than that before cold storage, and ethylene production and the SSC increased upon removal from cold storage. Fruits stored at RT reached the OTP on the 9th day, and the cold storage fruits reached the OTP three days earlier, whereas the stages in fruit maturity on the OTP with RT and after cold storage showed no significant differences.

Cold storage is an important method of maintaining fruit quality during postharvest storage. However, the fruits are subjected to chilling injury after long-term storage at LT. Chilling injury also induces aroma weakening, browning of the pericarp, and a reduction in shelf life (Zhou et al., 2014; Sheng et al., 2016; Wang et al., 2017). Previous studies have shown that cold storage reduces aroma production in peaches, tomatoes, and mangoes (Brian et al., 2015; Velu et al., 2017; Zhang et al., 2011). In the present study, RT was used as the control temperature, and LT was employed as the treatment temperature of ‘Nanguo’ pears to study the effect of LT storage on the expression of miRNAs that are related to fruit aroma volatile synthesis. The current study showed that 150 d of LT storage accelerates fruit ripening and senescence in ‘Nanguo’ pears. Furthermore, the aroma ester levels of the fruits were significantly lower during the OTP after cold storage than those at RT. These findings provide additional information on the mechanism underlying aroma weakening, which was previously associated with changes in the expression of regulatory genes related to aroma formation (Zhou et al., 2014; Shi et al., 2017).

Aroma volatiles are biosynthesized mainly through a sequence of enzyme reactions involving the lipoxygenase (LOX) pathway (Schwab et al., 2008). LOX is involved in the biosynthesis of 9-hydroperoxide and 13-hydroperoxide, which are derived from C6 and C9 aldehydes and catalyzed by HPL. These aldehydes are reduced to the corresponding alcohols by alcohol dehydrogenase (ADH) (EI Hadi et al., 2013). The availability of alcohols and acyl-CoA, and the properties of alcohol acyltransferases (AATs), determine the composition of volatile esters in fruits. AAT transfers acyl-CoA into the corresponding alcohol, and esters are generated by ester linkages (Tieman et al., 2006). *PuLOX1*, *PuLOX8*, *PuADH3*, and *PuAAT* may play important roles in aroma ester formation during pear ripening (Li et al., 2014). The roles of the *LOX*, *HPL*, *ADH*, and *AAT* genes in aroma weakening in ‘Nanguo’ pears during storage at RT after LT storage have been identified (Zhou et al., 2014; Shi et al., 2017). In the present study, the key genes *LOX2S* and *HPL* were significantly downregulated in fruits on the day they were removed from cold storage, and the *LOX2S*, *LOX1_5*, and *ADH1* genes were significantly downregulated in the fruits at the OTP after cold storage compared to those at the OTP at RT, and these genes showed no significant changes during aroma formation in fruits stored at RT. Moreover, the four key genes were predicted as the target genes of known significantly differentially expressed miRNAs in ‘Nanguo’ pears after cold storage.

In recent years, high-throughput sequencing technology has been widely used to identify miRNAs in plants, opening up a new area for research investigations on various molecular mechanisms. LT stress, a common abiotic stress that severely affects normal plant growth, development, and quality, has also received increasing attention, and significant progress has been made in understanding the role of miRNAs in response to LT stress (Zhang et al., 2009; Zhang et al., 2014). MiRNAs have been increasingly studied in response to LT stress in the past few years. However, no studies involving ‘Nanguo’ pears have been conducted to date. In the current study, to identify LT-responsive miRNAs related to aroma weakening in ‘Nanguo’ pears, four miRNA libraries of the fruit were established, which consisted on average of 10,878,552 (C0), 10,675,515 (COTP), 17,058,667 (LT0), and 12,601,065 (LTOTP) clean reads that were generated by high-throughput sequencing. Additionally, the differentially expressed miRNAs in fruits stored in the cold were identified, and mdm-miR172a-o, mdm-miR167b-g, mdm/ppe-miR395a-i, mdm-miR160a-e, mdm-miR7120a/b, mdm-miR159mdm-miR164b-f, and mdm-miR396b were most significantly differentially expressed in the fruits after LT storage. These low temperature-related miRNAs have also been identified in other plants (Chinnusamy et al., 2007; Josine et al., 2011; Karimi et al., 2016).

MiRNAs are post-transcriptional regulators of responses to abiotic stresses in plants and have been found to be multi-responsive to multi-environmental conditions individually or together with their target genes (Liu et al., 2008; Zhang et al., 2009; Lee et al., 2010). The miRNA families and their target genes in plants responding to LT stress include miR394, miR399, miR395, miR408, miR169, miR172, miR167, miR160, and miR159 (Cao et al., 2014; Xu et al., 2013; Karimi et al. 2016). MiR169 was differentially expressed by the NFYA gene, and the expression of miR160 and miR167 was regulated by auxin-response factors (ARFs) that were related to cold stress (Chen et al., 2012; Karimi et al. 2016). MiR156 and miR172 are the most important cold-responsive miRNAs (Xing et al., 2010), and the effect of miR156 and miR172 on LT has been reported in rice, wheat, and peach (Wang et al., 2014; Zhu et al., 2011). Although different species express different sets of miRNAs that respond to LT, there is a set of core miRNAs that is shared by most species. In the current study, miRNAs that mainly include mdm-miR172a-o, mdm-miR395a-i, mdm/ppe-miR399a, mdm-miR159a-c, mdm-miR160a-e, mdm-miR7120a/b, and mdm-miR535a were uniquely significantly differentially expressed during aroma formation in ‘Nanguo’ pears after cold storage, and the predicted target genes of these significantly differentially expressed miRNAs include the key aroma formation genes *LOX*, *HPL*, and *ADH*. Furthermore, we found that the expression of these target genes is related to these miRNAs. RT-PCR analysis indicated that mdm-miR172a-h, mdm-miR159a/b/c, mdm-miR160a-e, mdm-miR395a-i, mdm/ppe-miR399a, mdm/ppe-miR535a/b, and mdm-miR7120a/b are negatively regulated by the *LOX2S*, *LOX1_5*, *HPL*, and *ADH1* genes. Therefore, these miRNAs may be key factors in the aroma weakening of ‘Nanguo’ pears after LT storage.

In conclusion, the present study has identified differentially expressed miRNAs in response to LT during aroma formation in ‘Nanguo’ pears, which included mdm-miR172a-h, mdm-miR159a/b/c, mdm-miR160a-e, mdm-miR395a-i, mdm/ppe-miR399a, mdm/ppe-miR535a/b, and mdm-miR7120a/b. Furthermore, their predicted target genes play important roles in the aroma weakening of ‘Nanguo’ pears that were subjected to cold storage. The findings of the present study may be potentially utilized in future studies on the molecular mechanisms underlying the chilling injury of fruits subjected to cold storage. This study also provides valuable information for better understanding the roles of miRNAs in mediating aroma formation in fruits stored at LT.

## Acknowledgments

The National Natural Science Foundation of China (Grant number 31570687) supported this study.

